# *IPT9*, a cis-zeatin cytokinin biosynthesis gene, promotes root growth

**DOI:** 10.1101/2022.05.03.490520

**Authors:** Ioanna Antoniadi, Eduardo Mateo-Bonmatí, Markéta Pernisová, Federica Brunoni, Mariana Antoniadi, Mauricio Garcia-Atance Villalonga, Anita Ament, Michal Karády, Colin Turnbull, Karel Doležal, Ales Pěnčík, Karin Ljung, Ondřej Novák

## Abstract

Cytokinin and auxin are plant hormones that coordinate many aspects of plant development. Their interactions in plant underground growth are well established, occurring at the levels of metabolism, signaling, and transport. Unlike many plant hormone classes, cytokinins are represented by more than one active molecule. Multiple mutant lines, blocking specific parts of cytokinin biosynthetic pathways, have enabled research in plants with deficiencies in specific cytokinin-types. While most of these mutants have confirmed the impeding effect of cytokinin on root growth, the *ipt29* double mutant instead surprisingly exhibits reduced primary root length compared to wild type. This mutant is impaired in *cis*-zeatin (*c*Z) production, a cytokinin-type that had been considered inactive in the past. Here we have further investigated the intriguing *ipt29* root phenotype, opposite to known cytokinin functions, and the (bio)activity of *c*Z. Our data suggest that despite the *ipt29* short-root phenotype, *c*Z application has a negative impact on primary root growth and can activate a cytokinin response in the stele. Grafting experiments revealed that the root phenotype of *ipt29* depends on local signaling which does not relate to directly to cytokinin levels. Notably, *ipt29* displayed increased auxin levels in the root tissue. Moreover, analyses of the differential contributions of *ipt2* and *ipt9* to the *ipt29* short-root phenotype demonstrated that, despite its deficiency on *c*Z levels, *ipt2* does not show any root phenotype or auxin homeostasis variation while *ipt9* mutants were indistinguishable from *ipt29*. We conclude that IPT9 functions may go beyond *c*Z biosynthesis, directly or indirectly, implicating effects on auxin homeostasis and therefore influencing plant growth.

## INTRODUCTION

Plant roots are a highly powerful and dynamic part of the plant body that builds its underground architecture in search of water, anchorage and nourishment. Thus, the regulation of root growth and development is essential to plant prosperity and to their adaptability in changing environmental conditions. Cytokinins (CKs) have been long known inhibitors of root growth and development (1) and multiple mutants blocking CK biosynthesis, display enhanced primary root length compared to the wild type plants (2). In the past, *cis*-zeatin (*c*Z)-forms of cytokinin (CK) were considered less important than isopentenyl adenine (iP) and *trans*-zeatin (*t*Z)-forms due to their weaker responses in some bioassays (3,4) but also due to lack of research on them. More recently, it has been shown that *c*Z can be perceived by CK receptors in *Arabidopsis thaliana* (hereafter, Arabidopsis) and *Zea mays* (hereafter, maize; (5–8)) and that they are also bioactive in several assays (9). *c*Z-types were also detected as the main CK compound in developing seeds of chickpeas (*Cicer arietinum;* (10)), in all tissues of maize (11), in the flag leaves of rice (*Oryza sativa;* (12)) and during embryogenesis of pea (*Pisum sativum;* (13)). Additional evidence for *c*Z activity were presented when enzymes responsible for zeatin-O-glucosides production showed striking preference in *c*Z conjugation in maize (11). Similarly, the *c*Z-O-glucosyltransferases (*c*ZOGT1,2,3) identified in rice preferentially catalyze O-glucosylation of *c*Z-CKs than *t*Z-CKs (14). The suggestion that *c*Z has physiological effects on plant development was based on the phenotypes of *c*ZOGT overexpressor lines in rice, which displayed defects in crown root numbers, leaf senescence and shoot size (14). The direct impact of *c*Z activity in root elongation impairment was also exhibited *in tandem* with upregulation of CK response genes in rice (14). Salinity stress caused fast accumulation of *c*Z and the *c*Z precursor, *c*ZR (*cis*-zeatin riboside) in maize roots, while no change was observed in *t*Z levels (15). Similar increases in *c*Z-types were also observed following drought (16), heat (17) and biotic stress (18). In addition, a biological role for *c*Z-types in the regulation of xylem specification was recently reported (19). However, even though the double mutant of tRNA-AtIPTs, *ipt29*, had undetectable levels of all *c*Z-types, it displayed only chlorotic phenotype and was otherwise developmentally normal (2). Overall, *c*Z-types have been detected in more than 150 plant species and regardless their evolutionary complexity (9).

CK biosynthesis is catalyzed by 9 isopentenyl transferase enzymes (IPTs) in Arabidopsis (2). IPT1 and IPT3-8 are responsible for isopentenyl adenine (iP) and *t*Z compounds production (2). The biosynthesis of the latter ones requires an additional carboxylation step catalyzed by two cytochrome P450 enzymes (CYP735A1 and CYP735A2) (20,21). In parallel, IPT2 and IPT9 are responsible for *c*Z-type production following tRNA degradation (2). CK compounds can be categorized in two ways: (1) according to their side chain modifications (iP-, *t*Z- and *c*Z-compounds), and (2) to the changes in their adenine molecule during their metabolism (the precursor forms: phosphates and ribosides, the active free bases and the catabolite products: *N*- and *O*-glucosides (reviewed in (22)). The participation of *c*Z-compounds in CK homeostatic mechanism was demonstrated by the unanimous increased concentrations of *c*Z-types when *tZ*-types were deficient, in *ipt1 ipt3 ipt5 ipt7* (*ipt1357*) and *cyp735a1 cyp735a1a2* (*cypa1a2*) multiple mutants (20,23) and in *abcg14* (24,25). Interestingly, *c*Z-CK levels were increased in the mutants mentioned above only when *t*Z-types levels were reduced but not iP-types. The identification of *c*ZR as a major transport form of CK could also contribute to maintenance of CK homeostasis in the shoots (26).

The active CK molecules, free bases, are perceived by hybrid histidine kinases (HKs) in the CK-responsive cells and transcription is activated via a phospho-relay signaling cascade (27). The final step of this signaling network involves type-B nuclear RESPONSE REGULATOR (B-RR) proteins that mediate transcriptional activation by binding to promoters of immediate-early target genes via a conserved Myb-related DNA-binding domain. Global RR-B transcriptional activity and thus *in vivo* monitoring of these CK-dependent transcriptional responses, can be facilitated by the CK-responsive synthetic reporter *TCSn::GFP* expression in Arabidopsis plants (28). The signal output of this reporter line has also been shown to reflect CK content (29).

Although CK plays a pivotal role in root growth and development, several studies have shown that appropriate plant underground growth also depends on CK cross-talks with other phytohormones, such as indole-3-acetic acid (IAA), the main auxin. Derived from the amino acid L-Tryptophan (Trp), parallel biosynthetic and inactivation pathways converge to control IAA concentration (30). The main biosynthetic route, deaminates Trp to indole-3-pyruvic acid (IPyA) which is then decarboxylated to IAA. Although not completely clear, other routes exist connecting IAA biosynthesis with defensive compounds (glucosinolates) via indole acetaldoxime (IAOx). It has been hypothesized that IAOx, is then converted to indole acetonitrile (IAN) which is finally transformed into IAA. IAA levels are also controlled by redundant inactivation mechanisms including conjugation to sugars catalysed by different UGTs (42), GH3-driven conjugation to amino acids (43,44), and oxidation via DAO enzymes (45,46).

In this work, we monitored the primary root growth of three multiple mutant lines *ipt357, cypa1a2* and *ipt29*, which have impeded production of iP-, *t*Z- and *c*Z-type compounds, respectively. While *ipt357* and *cypa1a2* enhanced root growth is in agreement with the inhibitory effect of CKs on primary root growth, *ipt29* roots displayed severe retardation compared to wild type root length as also previously reported (19). We therefore investigated further the *ipt29* root phenotype, opposite to CK known functions, and the (bio)activity of *c*Z in the root tip. Our results showed that in spite of the defective growth of *ipt29* root, exogenous application of *c*Z triggers CK responses in the root vasculature and halts primary root growth. In parallel, grafting experiments indicated that *ipt29* root phenotype relies mainly on local signaling. Since this root signal was proven to be CK-independent, we examined auxin as a possible candidate, finding enhanced auxin levels the roots of *ipt29*. Finally, we analysed the differential contribution of *ipt2* and *ipt9* to the short-root phenotype. Surprisingly, even though there was a remarkable effect on *c*Z levels, *ipt2* root length and IAA levels were found normal while the *ipt9* single mutant showed a strong root phenotype and IAA levels indistinguishable from *ipt29*. No additional insertions in *ipt9* were found by genome-wide sequencing, and complementation tests and transgenic overexpression further confirmed the link between the short root phenotype and lesions in *IPT9*. Our data suggests that IPT9 bifunctionally works on *c*Z and IAA homeostasis and promotes root growth.

## MATERIALS AND METHODS

### Plant material, culture conditions and root phenotyping

Unless otherwise stated, all *Arabidopsis thaliana* plants studied in this work were homozygous for the mutations indicated. Single mutants *ipt2* (2), *ipt9-1* (2) multiple mutant *ipt357* (2), *cypa1a2* (20), *ipt29* (2), and the transgenic reporter line *TCSn::GFP* (28), *ARR5_pro_:GUS* (55), all in Col-0 background, were previously described. The Nottingham Arabidopsis Stock Centre provided seeds for the wild-type accession Col-0 (N1092), and *ipt9-2* (GABI_302F10; N428966).

The presence and position of all insertions were confirmed by PCR amplification using gene-specific primers, together with insertion-specific primers (Supplemental Table 1). In all experiments the seeds were surface sterilised with 20% (v/v) dilution of bleach for 5 min (2 × 2.5 min) and then rinsed five times with sterile water before sown under sterile conditions on Petri dishes containing half-strength Murashige and Skoog agar medium with 1% of sucrose. Stratification occurred at 4°C for 3 days and then plates were transferred to light at 22 ± 1 °C where the seedlings grew for 7 days under cool white fluorescent light (maximum irradiance 150 μmol m^-2^ s^-1^). For primary root length phenotyping the plates were scanned with Epson Perfection V600 Photo. Length quantifications were performed using FIJI software (56).

### Seedling treatments

Plants were grown in the above-described solid media supplemented with 100 nM of iP, *t*Z and *c*Z (Olchemim), respectively. After 7 days the plates were scanned and the root length was measured. For the root elongation assay, the seedlings were grown for 6 days in hormone-free media (as described in the previous section) prior to transfer to the media supplemented with 100 nM iP, *t*Z and *c*Z, under sterile conditions. The seedlings’ root tip positions were marked on the plates and they were returned to the growth chamber for 24h. After that, the plates were scanned and root elongation was measured.

### Histochemical staining

*ARR5_pro_:GUS* seedlings were stained in 0.1 M sodium phosphate buffer (pH 7.0) containing 0.1% X-GlcA sodium salt (Duchefa), 1 mM K_3_[Fe(CN)_6_], 1 mM K_4_[Fe(CN)_6_] and 0.05% Triton X-100 for 30 minutes at 37°C and were incubated overnight in 80% (vol/vol) ethanol at room temperature. Tissue clearing was conducted as previously described in (57).

### Microscopy

GFP expression patterns in 7-day-old *TCSn::GFP* seedlings was recorded using confocal laser scanning microscopy (Zeiss LSM800). The 488 nm laser line was employed for the GFP fluorescence detection, and emission was detected between 490 and 580 nm. Two tile scans were performed for root imaging. DIC microscopy was performed on an Olympus BX61 microscope equipped with ×10 and ×20 air objectives and a DP70 CCD camera.

### Auxin and Cytokinin measurements

Wild type and mutant plant roots were excised after 7 days of growth. The tissue was weighted and snap frozen in liquid nitrogen until hormone purification and analysis when frozen samples were thawed on ice.

For CKs analysis, samples (10 mg fresh weight) were homogenized and extracted in 0.5 ml of modified Bieleski buffer (60% MeOH, 10% HCOOH and 30% H_2_O) together with a cocktail of stable isotope-labelled internal standards used as a reference (0.25 pmol of CK bases, ribosides, N-glucosides, and 0.5 pmol of CK O-glucosides, nucleotides per sample added). CKs were purified using in-tip solid-phase microextraction based on the StageTips technology as described previously (58). Briefly, combined multi-StageTips (containing C18/SDB-RPSS/Cation-SR layers) were activated sequentially with 50 μl each of acetone, methanol, water, 50% (v/v) nitric acid and water (by centrifugation at 434×*g*, 15 min, 4 °C). After application of the sample (500 μl, 678×*g*, 30 min, 4 °C), the microcolumns were washed sequentially with 50 μl of water and methanol (525×*g*, 20 min, 4 °C), and elution of samples was performed with 50 μl of 0.5M NH4OH in 60% (v/v) methanol (525×*g*, 20 min, 4 °C). The eluates were then evaporated to dryness *in vacuo* and stored at −20 °C. The CK profile was then quantitatively analysed by multiple reaction monitoring using an ultra-high performance liquid chromatography-electrospray tandem mass spectrometry (UHPLC-MS/MS). Separation was performed on an Acquity UPLC i-Class System (Waters, Milford, MA, USA) equipped with an Acquity UPLC BEH Shield RP18 column (150×2.1 mm, 1.7 μm; Waters), and the effluent was introduced into the electrospray ion source of a triple quadrupole mass spectrometer Xevo TQ-S MS (Waters).

Analysis of endogenous IAA precursors and metabolites was performed using the method described in (59). Briefly, approx. 2.5 mg of roots or 10 mg of shoots were extracted in 50 mM phosphate buffer (pH 7.0) containing 0.1% sodium diethyldithiocarbamate and stable isotope-labelled internal standards. 200 μl portion of each extract was acidified with 1M HCl to pH 2.7 and purified by in-tip micro solid phase extraction (in-tip μSPE). For quantification of IPyA, the second 200 μl portion of the extract was derivatized by cysteamine (0.75 M, pH 8.2) for 15 minutes, acidified with 3M HCl to pH 2.7 and purified by in-tip μSPE. After evaporation under reduced pressure, samples were analyzed using HPLC system 1260 Infinity II (Agilent Technologies, Santa Clara, CA, USA) equipped with Kinetex C18 (50 mm x 2.1 mm, 1.7 μm; Phenomenex Torrance, CA, USA). The LC system was linked to 6495 Triple Quad mass spectrometer (Agilent Technologies).

CK and auxin concentrations were determined using MassLynx software (v4.2; Waters) and Mass Hunter software (version B.05.02; Agilent Technologies), respectively, using stable isotope dilution method. At least four independent biological replicates were performed, including two technical replicates of each.

### Grafting Experiments

Plants were grown vertically on 100 mm square plates containing 25 ml of 1/2 MS growth media at pH 5.7. Seeds (10 mg) were added to a 1.5ml Eppendorf tube with 1 ml of 70% ethanol and shaken at 21°C for 30 seconds. After washing with 1 ml of milliQ water, seeds were surface sterilised for 6 minutes in 1 ml of 1:10 diluted bleach, then washed six times with 1 ml of milliQ water. A 200 μl Gilson pipette was then used to suck up one seed at a time and transfer it on to plates, that were then kept at 4°C for 48 hours for stratification. The plates were then transferred to a controlled environment room at 23°C with a 16 hour photoperiod. After 5 days, plates were placed in a lit cupboard at 27°C. At 6 days post-stratification, simple hypocotyl grafting without a supporting collar was performed according to Turnbull et al. 2002. The non-grafted, self-grafted and trans-grafted seedlings were placed again at 27°C for another 4 days, then transferred back to 23°C. The plants were checked every day for evidence of contamination and to eliminate any adventitious roots growing above the grafting incision. This was carried out causing minimum disturbance to allow for graft union formation (60). Eight days post grafting the plates were scanned and the root length was measured in all successful grafts.

### Transgene complementation

To construct the *35S_pro_:IPT9*, the At5g20040 transcription unit was amplified from Col-0 cDNA using the Q5 High-Fidelity DNA Polymerase (NEB), as recommended by the manufacturer using the oligonucleotides attB_IPT9_F and attB_IPT9_R, that contained attB sites at their 5’ ends (Supplemental Table 1). The PCR product obtained were purified using the Monarch DNA Gel Extraction Kit (NEB) and cloned into the pDONR207 using a BP Clonase II Kit (Thermo Fisher). Chemically competent *Escherichia coli* DH5α cells were transformed by the heat-shock method (Dagert and Ehrlich 1979), and the structural integrity of the inserts carried by transformants was verified by Sanger sequencing. The insert cloned in the pDONR207 was subcloned into the pMDC32 Gateway-compatible destination vector (61) via an LR Clonase II (Thermo Fisher) reaction. The transgene was mobilized into *Agrobacterium tumefaciens* GV3101 (C58C1 Rif^R^) cells and those were used to transform Col-0, *ipt9-1* and *ipt9-2* plants by the floral dipping (62). T_1_ transgenic plants were selected on plates supplemented with 15 mg/L Hygromycin B (Duchefa).

### Genome-wide identification of additional T-DNA insertions in KG7770

We sequenced the *ipt9-1* (KG7770) genome and followed a tagged-sequence strategy to map the potential additional insertions as described in (44). Briefly, 8.6 μg of nuclear-enriched DNA was purified from 0.5 g of *ipt9-1* seedlings as previously described in (63). Whole-genome sequencing of the sample was performed at BGI Hong Kong using a BGISEQ-500 sequencing platform. 28.93 million 150-bp-long reads were obtained, reaching a 32X genome depth. Trimmed fastq files were used to map the position of the insertion using Easymap software (64). Raw reads were deposited in Short Read Archive with the code SRX14238175.

## RESULTS

### Cytokinin deficient mutants display different primary root phenotypes

CK biosynthesis pathway can be blocked, in principle, in three different levels targeted by the mutants *ipt357* (2), *cypa1a2* (20) and *ipt29* (2). These three mutant lines are known for impaired iP-, *t*Z and *c*Z- biosynthesis, respectively (Figure 1A). CK’s inhibitory effect in root growth has been well established (1) and most CK deficient and insensitive mutants display longer primary roots accordingly (65). Here, we have grown *ipt357, cypa1a2* and *ipt29* mutants for seven days and assessed their primary root growth. The mutants *ipt357* and *cypa1a2* displayed longer primary root phenotypes compared to Col-0 (Figure 1B-C) while *ipt29*, impaired in *c*Z production, exhibited a severely reduced root growth (Figure 1B-C). While this phenotype of *ipt29* has been previously described by other authors (19), it remains enigmatic. Therefore, here we aimed to elucidate the *ipt29* short-root mutant phenotype by using the *ipt357* and *cypa1a2* mutants as controls.

**Figure 1.**
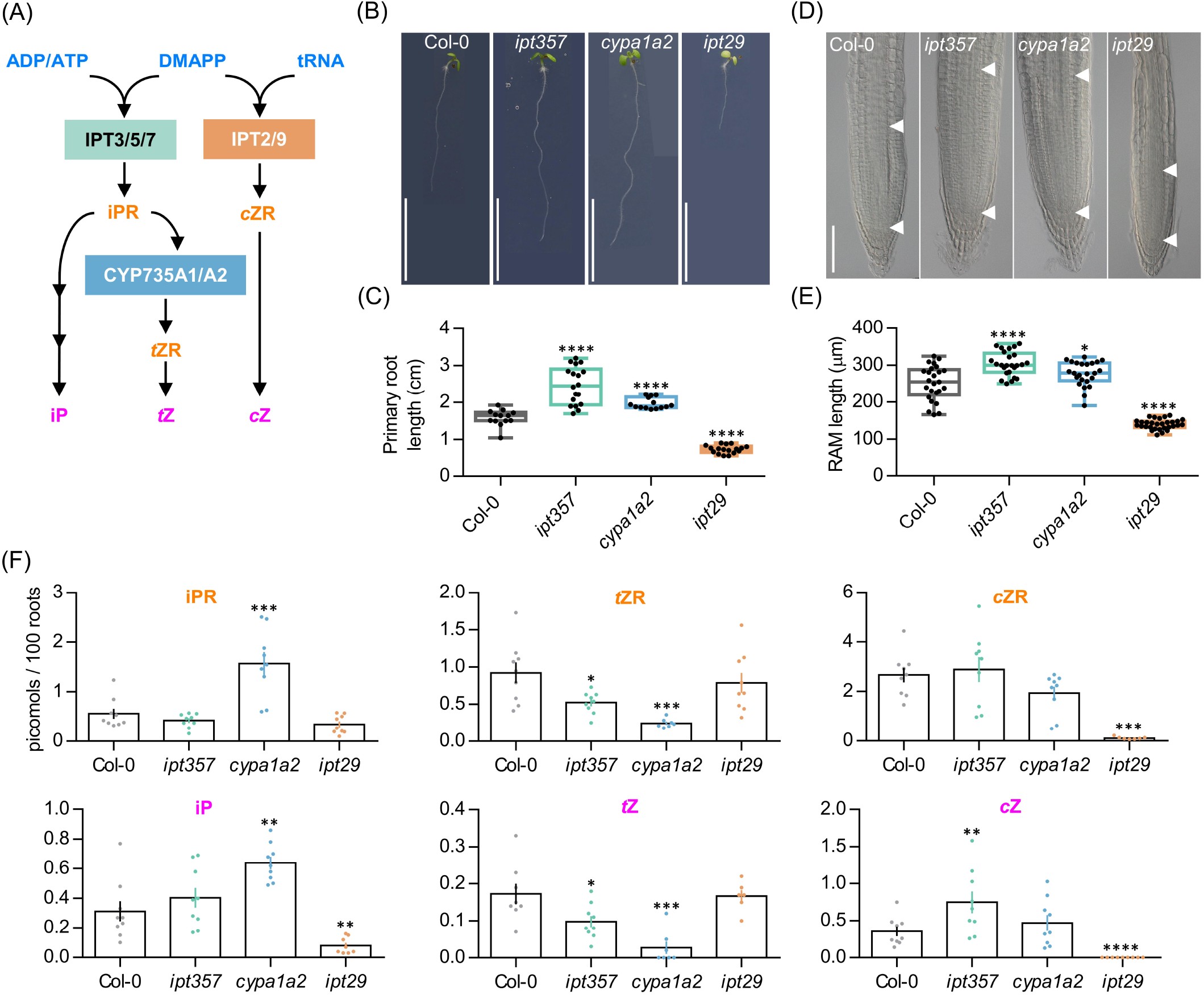
Mutants in different cytokinin biosynthetic genes show differential effects on primary root length and meristem size. (A) Simplified version of the parallel cytokinin biosynthesis pathways. (B, C) Primary root phenotype and length quantification of the wild-type Col-0, the triple mutant *ipt357*, and the double mutants *cyp753a1 cyp753a2* (*cypa1a2*) and *ipt29*. (D, E) Root meristem phenotype and size quantification of the assorted genotypes. (F) Quantification of some biologically active cytokinins in the assorted genotypes expressed as picomoles per 100 roots. Pictures were taken 7 days after stratification (das). Scale bars indicate (B) 1 cm, and (D) 100 μm. Asterisks indicate values significantly different from Col-0 in a (C, E) Student’ *t* test and (D) Mann-Whitney *U* test (*p<0.05, **p<0.01, ***p<0.001, ****p<0.0001; n≥ (C) 13, (E) 24, (F) 6). ADP/ATP, adenosine di/triphosphate; DMAPP, dimethylallyl diphosphate; tRNA, transferTransfer ribonucleic acid; IPT, isopentenyltransferase; iPR, isopentenyladenosine; *c*ZR, *cis*-zeatin riboside; *t*ZR, *trans*-zeatin riboside; iP, isopentenyladenine; *t*Z, *trans*-zeatin; *c*Z, *cis*-zeatin.

The primary root defect observed in *ipt29* mutant was consistent with shorter meristematic root zone size compared to Col-0 (Figure 1D-E) and its CK metabolic profile showed severely lower levels of *c*ZR and *c*Z but also of iP compared to wild type plants (Figure 1F). Both the *ipt357* and the *cypa1a2* mutants displayed lower contents for *t*Z riboside and *t*Z compared to Col-0, while iPR and iP levels were higher only in *cypa1a2* mutant (Figure 1F).

### *ipt29* short-root phenotype is cytokinin independent and is controlled by local signals

To assess whether exogenous CK supply could rescue the defective root phenotypes of these mutant lines, *ipt357, cypa1a2* and *ipt29* were grown for seven days in media supplemented with 100 nM iP, *t*Z or *c*Z. All three CK compounds inhibited root growth in all genotypes (Figure 2A). The inhibitory effect on root growth was similar for *t*Z and iP and lower for *c*Z both in the wild-type and all the mutant lines (Figure 2A-B). While the longer root phenotypes of *ipt357* and *cypa1a2* observed compared to Col-0 in mock conditions were inverted in response to CK treatments, the short root phenotype of *ipt29* still remained (Figure 2A). In fact, 100 nM cZ could fully rescue the *ipt357* and *cypa1a2* root phenotypes but not that of *ipt29* (Figure 2A). Similar results were observed in root growth elongation rate assays in which the same genotypes were grown for six days in MS media and then transferred for one day to media supplemented with 100 nM iP, *t*Z or *c*Z (Supplementary Figure 1A-B). None of these three active CKs could rescue the short-root phenotype of *ipt29* (Figure 2A), nor any of the *c*Z concentrations applied in the dose-response assay shown in Figure 2D.

**Figure 2.**
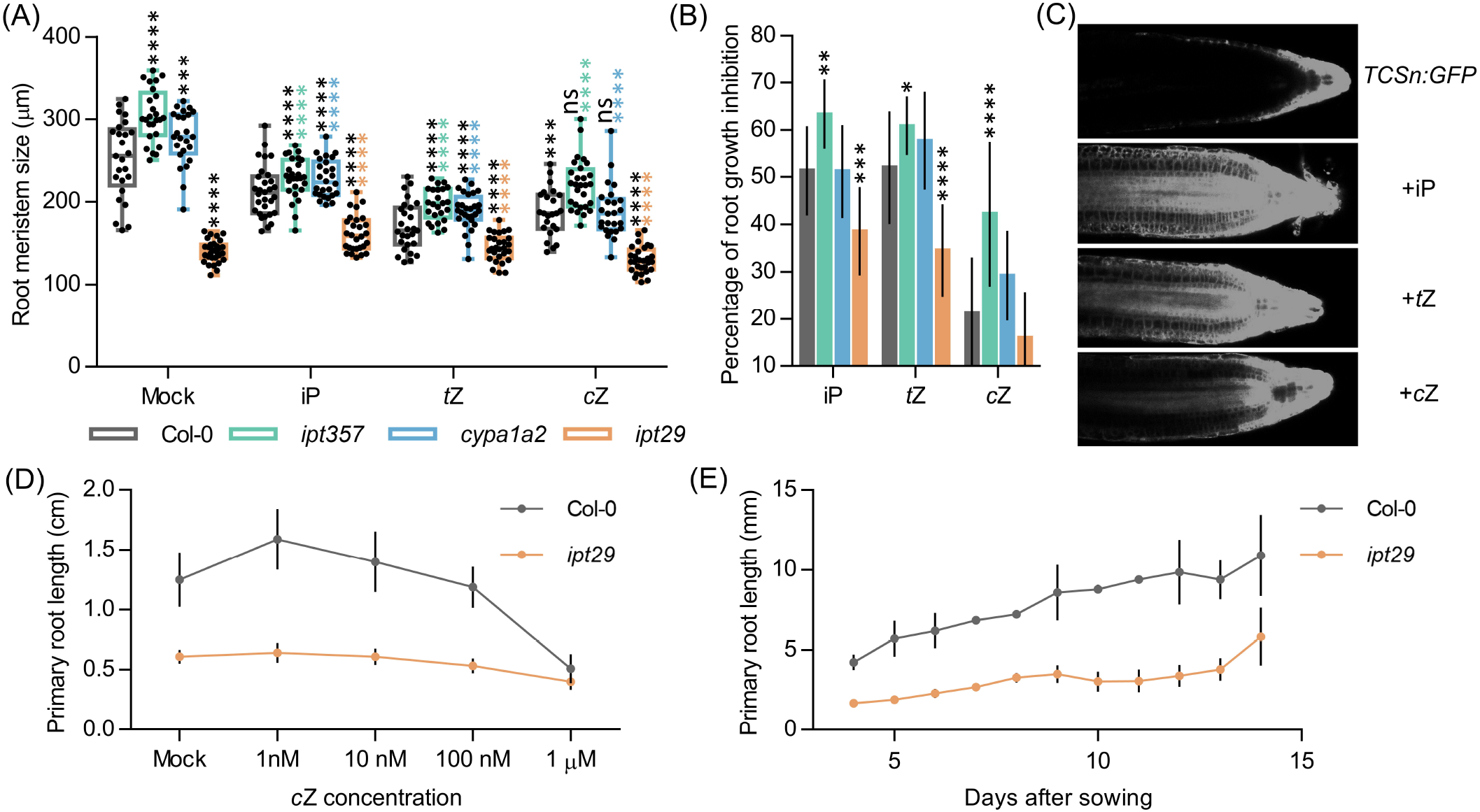
Mutants in different cytokinin biosynthetic genes show differential effects on primary root length and meristem size. (A) Primary root length of the wild-type Col-0, and the *ipt357, cypa1a2*, and *ipt29* multiple mutants grown for 7 days in media supplemented with 100 nM of iP, *t*Z and *c*Z. Black asterisks indicate values significantly different from Col-0 mock treatment and colour asterisks indicate significant differences from the corresponding mock genotype in a One-way ANOVA test (***p<0.001, ****p<0.0001; n ≥ 13). (B) Root growth inhibitory effects of iP, *t*Z and *c*Z on the genotypes mentioned in (A). (C) Cytokinin signalling reporter TCSn response to different cytokinin treatments. (D) *c*Z dose-response treatment of Col-0 and the ipt29 double mutant. Primary root length after 7 days growing in media supplemented with different *c*Z concentrations. Dots indicate average ± SD of at least 20 seedlings (E).

Induction of CK responses following 100 nM iP, *t*Z and *c*Z treatment was assessed with fluorescence imaging of root tips from the CK response reporter line *TCSn:GFP*. All three CKs were able to induce CK signaling when seedlings were grown on treated media (Figure 2C) or when seedlings were transferred for 24 h to media containing the respective treatment (Supplementary Figure 1C). Both the *TCSn:GFP* and the transcriptional fusion of *ARABIDOPSIS THALIANA RESPONSE REGULATOR5* (*ARR5*) promoter to the β-glucuronidase (*ARR5_pro_:GUS*) lines (55) displayed increased intensity of CK signaling in response to 24 h of different CK types, in the order: iP >*t*Z >*c*Z (Supplementary Figure 1C, D). Finally, comparing *ipt29* root phenotype with Col-0 when plants were grown for 14 days confirmed that the root remained significantly shorter in the mutant even after a longer growth period (Figure 2E).

The results obtained so far pinpointed that the *c*Z deficiency of the *ipt29* mutant most likely was not linked with the short-root phenotype, and therefore another non-CK signal could be the explanation. As a first step, we examined whether this signal was shoot-derived or local. To answer this, reciprocal grafts were performed between scions and rootstocks derived from Col-0 and *ipt29* genotypes, and grafts of the same genotype were used as controls, as shown in Figure 3A.

**Figure 3.**
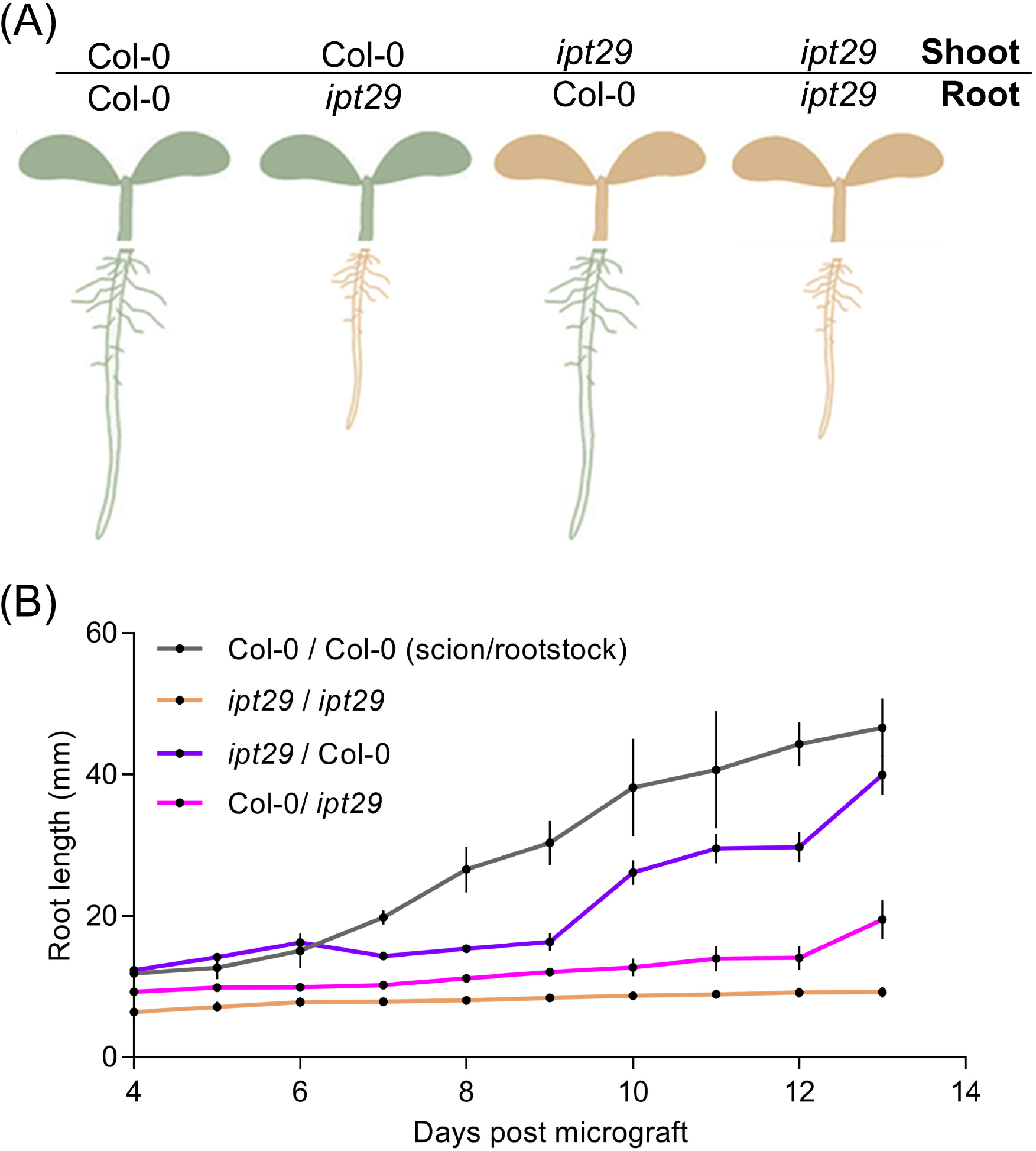
*ipt29* phenotype is mainly controlled by local signals. (A) Schematics representing grafting experiments performed between Col-0 and *ipt29* shoot and root. Col-0 and *ipt29* were self-grafted as controls. (B) Primary root growth of grafted seedlings. Dots indicate the average ± s.e.m. of the primary root length (n=24) of grafted and self-grafted seedlings of the assorted genotypes.

The grafts were performed when the plants were four-day-old and thereafter their root growth was recorded daily. As shown in Figure 3B, the control grafts Col-0/Col-0 (scion/rootstock) and *ipt29/ipt29* had the previously observed difference in primary root length (Figure 1B-C). The reciprocal grafts *ipt29*/Col-0 and Col-0/*ipt29* both displayed shorter roots compared to the wild type control graft. Since the Col-0 scion was not sufficient to restore the short-root phenotype of the *ipt29* rootstock (Figure 3B), we concluded that the signal controlling the phenotype is not shoot-derived and it is likely locally generated in root tissues.

### A strong link was observed between auxin metabolism and the *ipt29* short-root phenotype

Our results so far indicated that the shorter root phenotype of the *ipt29* mutant is not CK-dependent and is controlled locally in the root tissue. Another signal that often interacts with CK to regulate root growth and can act locally is auxin (66,67). Therefore, we carried out a detailed profiling of auxin and related metabolites, analysing separately shoot and root tissues for the active form IAA as well as several precursors such as anthranilate (ANT), Trp, IPyA, and IAN, and inactive forms like indole-3-acetyl glutamate (IAA-Glu), 2-oxindole-3-acetic acid (oxIAA) or 2-oxoindole-3-acetyl-1-O-β-D-glucose (oxIAA-glc) (Figure 4A, B). Both in shoot and root tissues, IAA levels were significantly higher in *ipt29* than Col-0, this difference being more acute in root tissues. In shoots, IAA levels were slightly higher in *ipt29* compared to wild type, while all the inactive forms analysed were depleted. Regarding the precursors, ANT levels were reduced in *ipt29* while the others were indistinguishable from Col-0 (Figure 4A). In roots, IAA metabolism was more strikingly affected with higher levels of IAA inactivated catabolites, while among the precursors only Trp levels were significantly higher in *ipt29* than Col-0 (Figure 4B). The chlorotic phenotype of *ipt29* leaves suggests some type of chloroplast misfunction, pointing to a potential defect on the IAA precursor Trp. Our results, however, discard any Trp deficiency and rather discovered a clear link between the *ipt29* mutation and higher IAA levels. Overall effects of auxin on root growth are very concentration dependent: low concentrations of IAA normally promote growth, but altering IAA homeostasis and enhancing IAA concentrations results in growth inhibition, potentially explaining the *ipt29* root phenotype.

**Figure 4.**
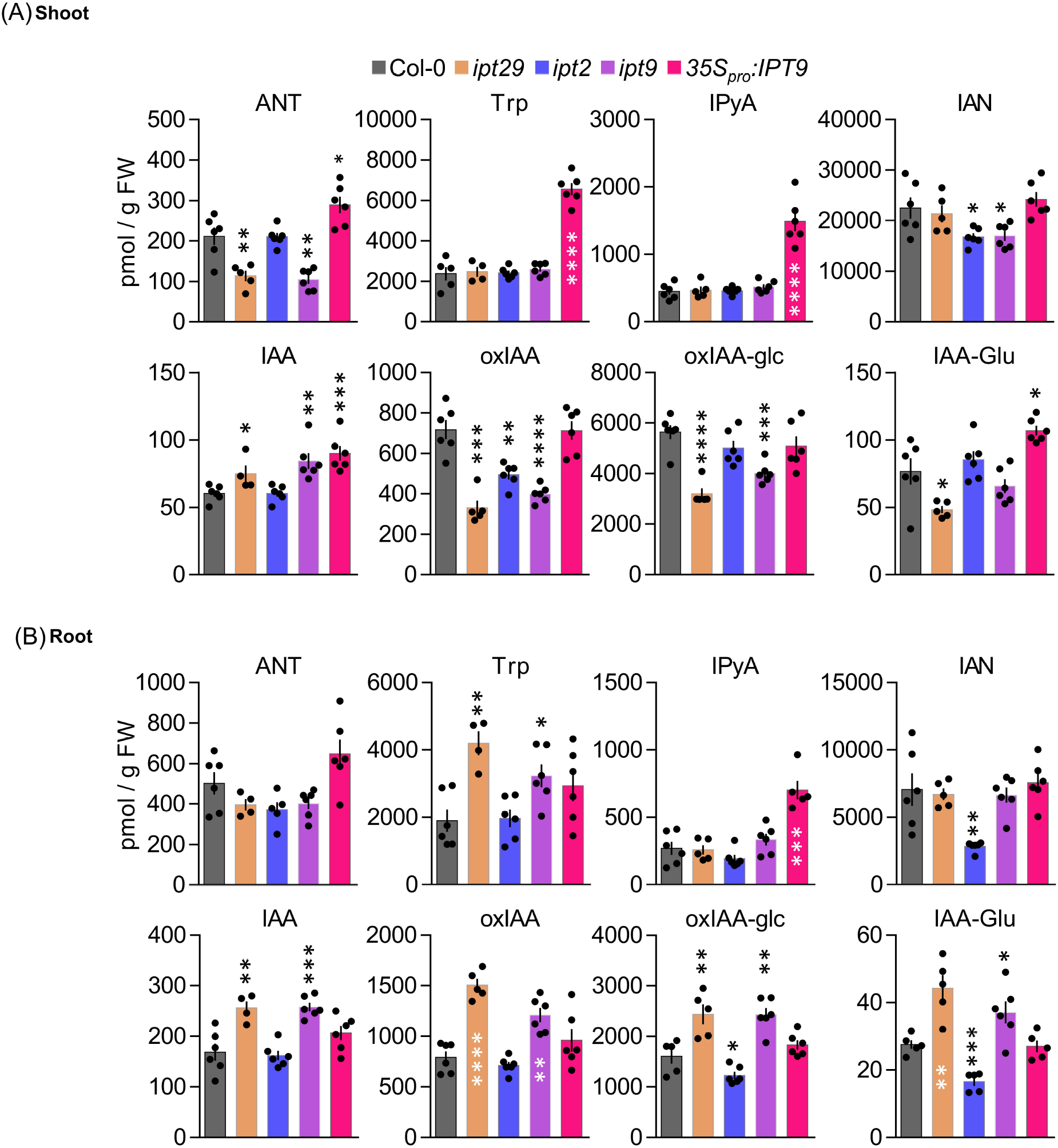
Shoot and root profiling of auxin metabolites in cZ biosynthetic mutants. Anthranilate (ANT), Tryptophan (Trp), indole-3-pyruvic acid (IPyA), indole-3-acetonitrile (IAN), indole-3-acetic acid (IAA), 2-oxindole-3-acetic acid (oxIAA), 2-oxoindole-3-acetyl-1-O-β-D-acetic acid glucose (oxIAA-glc), and indole-3-acetyl glutamateacetic acid glutamic (IAA-Glu) levels were quantified in (A) aerial tissues and (B) roots of Col-0, ipt29, ipt2, ipt9, 35S_pro_:IPT9 7-days-old seedlings. Concentrations are expressed in picomols per g of fresh weight. Error bars indicate standard error and asterisks indicate values significantly different from Col-0 in a Student’s t test (* p <0.05, ** p <0.01, *** p <0.001, **** p <0.0001; n ≥ 4).

### IPT9 is solely responsible for the *c*Z-independent *ipt29* phenotype

To ascertain whether there is a differential contribution to the *ipt29* phenotype from either of the two paralog genes, we analysed the primary root phenotype of both single mutants. Surprisingly, while *ipt2* single mutant root phenotype was indistinguishable from the wild type, *ipt9* roots were as short as those of the double *ipt29* (Figure 5A-B). We then wondered if levels of *c*ZR, *c*Z and IAA could explain these differential phenotypic contributions in the single mutants. However, concentrations of *c*ZR and *c*Z were much more reduced in *ipt2* than in *ipt9* roots, further confirming the CK-independence of the *ipt29* short root phenotype (Figure 5C). In line with the link found between the *ipt29* short root phenotype and IAA metabolism, *ipt9* showed higher IAA levels both in shoot and root while *ipt2* IAA levels were indistinguishable from Col-0 (Figure 4A, B).

**Figure 5.**
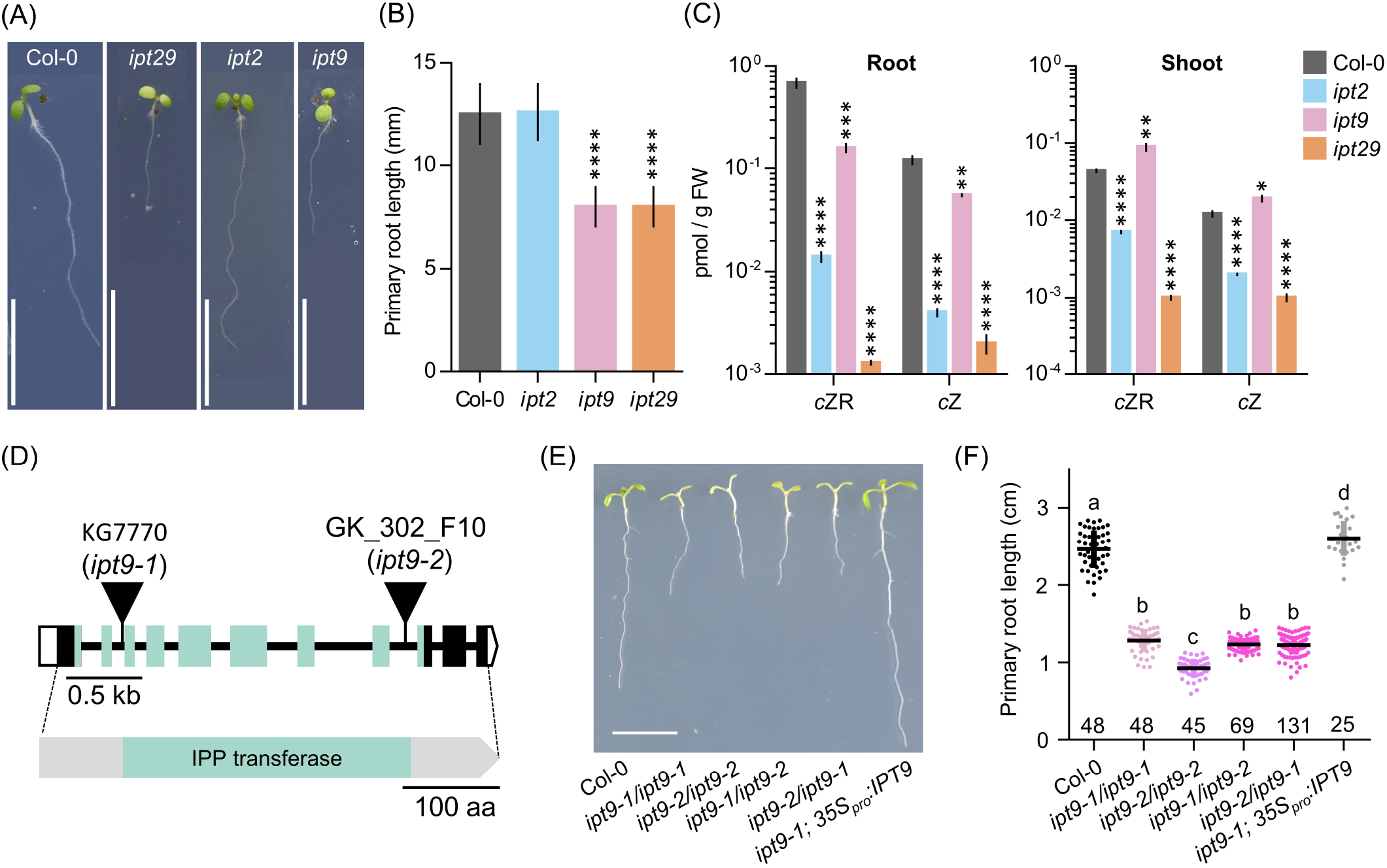
Lesions in *IPT9* are solely responsible of the *ipt29* phenotype and are *c*Z-independent. (A, B) Primary root phenotype (A) and length quantification (B) of Col-0, *ipt2, ipt9*, and *ipt29*. Asterisks indicate values significantly different from Col-0 in a Student’s t test (* p <0.05, ** p <0.01, *** p <0.001, **** p <0.0001; n=5). (C) *c*ZR and *c*Z quantification of shoot and root of Col-0, *ipt2, ipt9*, and *ipt29*. (D) Structure of the *IPT9* gene and protein with the nature and positions of the *ipt9* mutations indicated. Boxes and lines represent exons and introns, respectively. Open and coloured boxes represent untranslated and translated regions, respectively. Triangles indicate T-DNA insertions in *ipt9-1* and *ipt9-2* mutants. Greenish shadowed regions represent exons encoding and region of the protein corresponding to the tRNA delta(2)-isopentenylpyrophosphate transferase (IPP; pfam01715) domain. (E, F) Primary root phenotype (E) and length quantification (F) of Col-0, homozygous *ipt9-1*, and *ipt9-2*, trans-heterozygous *ipt9-1/ipt9-2*, and *ipt9-2/ipt9-1*, and transgenic plants in the *ipt9-1* background expressing *IPT9* under the constitutive 35S promoter. Different letters indicate values significantly different (p < 0.05; n of each population is indicated above the genotype) in a Tukey’s post-doc test. Pictures were taken (A, E) at 7 days. Scale bars indicate (A, E) 1 cm, (D) 0.5 kb and 100 amino acids.

Since all our analysis have been performed using previously reported T-DNA insertional mutants and it is well established that the average number of T-DNAs in insertional lines is greater than two (68), we wondered if an additional insertional event could explain the defects we observe in *ipt9*. To rule out this possibility, we sequenced the *ipt9* genome and followed a tagged-sequencing strategy to map the position of all the insertional events. We confirmed the presence of an insertion only in At5g20040 (*IPT9;* Supplemental Figure 2A). We also obtained a second insertional allele for *IPT9*, the line GK_302_F10 from the GABI-KAT collection. Therefore, we re-named the already published allele of *IPT9* (KG7770) as *ipt9-1*, and the GABI-KAT line as *ipt9-2*. The insertions interrupt the second (*ipt9-1*) and the eighth (*ipt9-2*) intron of the *IPT9* gene, both affecting the sequence coding for the tRNA delta(2)-isopentenylpyrophosphate transferase protein domain (IPP; pfam01715; Figure 5D). We then performed a phenotypic complementation assay with both alleles. All homozygous and trans-heterozygous plants combining *ipt9* mutant alleles showed significantly shorter roots than Col-0, thus confirming the direct relationship between *IPT9* misfunction and the *ipt9* and *ipt29* short root phenotype (Figure 5E-F), among other phenotypic traits such as chlorotic leaves (Supplementary Figure 2B-D) and shorter stems (Supplementary Figure 2E-G). We next transferred a transgene containing a wild-type version of *IPT9* driven by cauliflower mosaic virus 35S promoter to both *ipt9-1* and *ipt9-2* mutant lines (Figure 5E, F, Supplementary Figure 3 A-C). Detailed analysis of multiple independent transgenic families (selected from different T_0_ families) showed that not only in most of the transgenics the root phenotype was restored to wild-type levels but also that in some families, roots were even longer than Col-0 (Figure 5E, F, Supplementary Figure 3 A-C). To remove potential interactions between the wild type and the mutant version of IPT9 in any mutant background, we also introduced the transgene into Col-0 plants. Again, some families, 6 out of 10 analysed in detail, showed enhanced root growth compared to non-transgenic Col-0 (Supplemental Figure 3D, E). To explore the IAA metabolic landscape of these plants overexpressing *IPT9*, we performed auxin profiling of shoots and roots (Figure 4A, B). While in shoots the effects of this transgene increased the IAA levels even further, IAA levels in roots were the same as Col-0. Other differential shoot/root effects of the transgene such us the increased levels of ANT, Trp and IAA-Glu found in shoots were not mimicked in roots (Figure 4A,B). IPyA was the only exception, with higher levels in both tissues. In conclusion, our allelic and transgenic complementation combined with phenotypic and hormonal analyses demonstrated that mutations in the *IPT9* are wholly responsible for the *ipt29* short root phenotype.

## DISCUSSION

CKs are important for plant development as they are regulating multiple plant functions. In contrast with other plant hormones, such as auxins or brassinosteroids, CKs have more than one active molecule. These are the nucleobases iP, *t*Z, *c*Z and dihydrozeatin. They can bind to CK receptors and initiate the corresponding hormonal responses affecting various physiological aspects of plant growth such as root growth, contributing thus to plant development and adaptation. These four molecules have different affinities to their receptors (6–8,69), they can be produced in same but also in spatially different locations within the plant tissues (19–21,70–80) and they are degraded and conjugated at different rates (9,81,82). It is thus interesting to understand why this hormone has four, instead of one, active molecules and whether these could possibly have different effects in specific plant functions, such as root growth.

### The *ipt29* double mutant has a unique primary root phenotype

The triple and double mutants, *ipt357, cypa1a2* and *ipt29*, are inhibited in different parts of the CK iP, *t*Z and *c*Z biosynthesis pathways, respectively (Figure 1A). Previous studies have shown that *ipt357* has a longer primary root compared to wild type, supporting the inhibitory effect of CK on root growth (1) and our results show that this is also the case for the *cypa1a2* double mutant. Interestingly *ipt29*, specifically blocked *c*Z production, has shorter roots compared to wild type plants.

This intriguing phenotype that could potentially suggest that *c*Z has an opposite effect in root growth compared to the other active CK compounds, triggered not only our interest but also the one of Kollmer *et al* (19). In their work studying the *CKX7* gene, they found that CKX7 enzyme preferred *c*Z compounds as substrate compared to other CKs (9,19) and that plants that overexpressed *CKX7* had severely retarded root growth compared to wild type, similarly to *ipt29* root phenotype (19). Their results showed that *c*Z-CKs play an important, yet not exclusive, role in vascular differentiation as other CK-types could affect this process. However, the question of why the *ipt29* mutant has a shorter root phenotype remained unanswered.

Our results confirmed the defective primary root phenotype of *ipt29* in comparison with Col-0 and the other adenylate-*IPT* mutants that had longer primary roots (Figure 1B-C). The observed phenotypes were in accordance with shorter and longer meristematic zone of the mutants’ roots, respectively (Figure 1D-E). The correspondence of root meristem size and primary root length of *ipt29* mutant is in agreement with previous findings (19).

Initially, we compared the CK profiles between 7-day-old Col-0 and the CK-type deficient mutants. The results overall confirmed the expected reduced CK-types content in the respective mutant. In addition, the higher concentration of *t*Z and *t*ZR in *ipt357* and the reciprocal effect of higher CK iP-type and lower *t*Z-type levels in *cypa1a2* mutant roots, suggested that in young Arabidopsis roots most of the iPMRP produced by IPT action is converted into *t*ZRMP for *t*Z-rypes production. This supports the importance of *t*Z in Arabidopsis young roots in agreement with this compound’s prevalence in CK responsive cells of the same plant and tissue (29). In *ipt29* mutant roots, *c*Z-CKs concentrations were severely reduced as previously shown in older plants (2,19). Based on the reduced concentration of iP in *ipt29* and the increased *c*Z levels in *ipt357* mutants, an enigmatic hypothesis could be a potential enzymatic connection between iP and *c*Z biosynthesis pathways.

### *c*Z deficiency is not linked with the *ipt29* short-root phenotype

All active CK molecules (iP, *t*Z, *c*Z) were able to suppress the increased root growth (Figure 2A-B) and growth rate (Sup. Fig 1A-B) phenotypes of *ipt357* and *cypa1a2*, indicating that the longer root phenotypes in these mutants are caused by CK deficiency. These experiments (Figure 2A, B and Supplementary Figure 1A-B) also revealed that *c*Z has the weakest effect on root length and growth rate compared to the other two active CKs applied (*t*Z~iP>>*c*Z), implying that cZ could have a less important role in root growth regulation compared to the other two compounds. Although actual embryos have not been examined, embryonic effects seem unlikely to be the cause of the *ipt29* phenotype because of the following results. Firstly, the seeds of *ipt29* germinated at the same time as Col-0. Secondly, *ipt29* mutants grown for the longer period of 14 days displayed the same retarded root growth compared to the wild type as the 7-day-old seedlings (Figure 2E). Finally, the *ipt29* root defects could be also attributed to the lower root growth rate of this mutant (Supplementary Figure 1A-B) which is a post-embryonic process. Overall, since no applied concentration of *c*Z (Figure 2D) and no other CK treatment (Figure 2A) could alter the retarded root growth or elongation of *ipt29* mutant, it can be concluded that inhibition of root growth in *ipt29* mutant is a CK-independent phenotype.

In fact, when *c*Z was applied exogenously the root phenotype of *ipt357* and *cypa1a2* could be fully rescued (Figure 2A and Supplementary Figure 1A-B). This supports that *c*Z acts as an inhibitor of root growth, like the other active CKs, possibly following binding of *c*Z to AHK CK receptors and thus activating downstream signaling. Indeed, *c*Z treatment was able to activate the CK response reporter line, *TCSn:GFP* (28), although in lower intensity compared to iP and *t*Z (Figure 2C and Supplementary Figure 1C). It has been previously shown that CK receptors differ in their preference of CK isoforms (6,83,84) and Lomin et al, summarized in their review data supporting that *c*Z has lower affinity to AHK receptors compared to iP and *t*Z in both Arabidopsis and maize (85).

An alternative scenario for the *c*Z-driven responses mentioned above that cannot be excluded is that *c*Z, at least partially, converts to *t*Z via potential isomerization by zeatin cis-/trans-isomerase as shown in maize cultures. When these were incubated for 20 min with *c*Z, about one tenth of the compound was converted to *t*Z (8). Another explanation in our case could be that *c*Z conversion into *t*Z occurs via the hydroxylation of N6-isopentenyl side chain or other unknown reaction. In parallel, feeding experiments on Arabidopsis protoplasts and maize cultured cells with labelled *c*Z indicated to a metabolic route from *c*Z not only to respective conjugates but also back to its precursor forms (8,29). However, in such feeding experiments on rice seedlings, tobacco cells, oat leaves and Arabidopsis protoplasts no isomerization was observed between *t*Z- and *c*Z-ribosides (9,14,86). Likewise, lack of isomerization from *t*Z to *c*Z was inferred from absence of detectable *c*Z-types in the *ipt29* double mutant (2). The above studies suggest that *c*Z-type levels are pivotally controlled by *de novo c*Z-biosynthesis through the tRNA pathway. Lack of *cis-trans* isomerization also indicates that the high levels of *c*ZR being transported through the plant body (26) are more likely to have a biological role directly as *c*ZR or *c*Z rather than contributing to the *t*Z-CK pools of the sink.

### The *ipt29* mutant phenotype is governed by local signals in the root

Previous grafting experiments have revealed that CKs can act not only as local signals but also as distal ones. Interestingly, *c*Z-types and more specifically *c*ZR are prevalent compounds in phloem and xylem sap of Arabidopsis (26) although this compound seems to be almost inactive when tested *in vitro* receptor binding assays in Arabidopsis and maize (7,8). In agreement, *c*ZR was unable to initiate ARR5-mediated CK response (7). On the other hand, *c*ZR had strong effect in tobacco callus growth and oat chlorophyll retention bioassays (9). This indicates that *c*ZR can have great activity in bioassays probably after its conversion to the bioactive *c*Z. Transport of cZ CKs in inactive forms such as *c*ZR could provide a further regulatory level to control extent of CK responses.

Shoot-born CKs can act as an inhibitory signal of nodule formation on *Lotus japonicus* roots (87). The defective phenotype of *ipt1357* quadrupole mutant scion and rootstock could be restored by Col-0 rootstock and scion in reciprocal grafting experiments (23). Also, *cypa1a2* shoot phenotype was rescued when the mutant scion was merged with wild type rootstock (20). To verify that *ipt29* sort root phenotype is unrelated to endogenously transported CKs, grafting experiments were performed (Figure 3). Shoot-derived CKs from Col-0 scion were unable to restore the *ipt29* rootstock phenotype (Figure 3B) suggesting that the signal controlling this phenotype is a local signal in the root and according to our results it is not CK-dependent.

### Link to auxin metabolism and differential contribution of *ipt2* and *ipt9* to the short root phenotype

Auxin and CKs have been previously shown to interact at many levels such as biosynthesis, signalling and transport controlling several developmental processes including lateral root initiation, vascular development and meristem size and maintenance (88). Examples of CK-auxin crosstalk include the SUPPRESSOR OF HYPOCOTYL2 (SHY2), which controls root meristem activity by balancing auxin and CK responses. While the SKP1-CULLIN1-F-BOX (SCF)–TRANSPORT INHIBITOR RESISTANT1 (SCF^TIR1^) complex enables the auxin-dependent degradation of SHY2 (47,48), B-RRs-regulated CK signaling in the root meristem transition zone, directly activates *SHY2* transcription. SHY2 then negatively affects auxin efflux and response. In addition, SHY2 also induces CK biosynthesis by upregulation of *IPT5* (49,50) Interaction of the two hormones also control the activity of the quiescent center (QC). In fact, ARR1-dependent CK signaling causes downregulation of the auxin influx carrier *LIKE AUXIN RESISTANT2* (*LAX2*), leading to attenuation of auxin response and division in QC cells (51). In the QC, the transcription factor SCARECROW (SCR) suppresses ARR1 and affects auxin biosynthesis (52). Auxin also antagonizes CK by direct transcriptional activation of *ARR7* and *ARR15* that are repressors of CK signaling (53). A domain of high-auxin signaling in the xylem cells and a domain of high-CK signaling in the procambium and phloem cell lineages is described to control vascular tissue patterning. Very recently, auxin was found to activate the TARGET OF MONOPTEROS 5 and LONESOME HIGHWAY (TMO5/LHW) heterodimer complex that serves as central organizer for vascular development and patterning in the root apical meristem. The TMO5/LHW module, in turn, directly controls SHORTROOT (SHR) to balance CK levels (54).

Auxin was thus a good candidate signal to assess as a potential explanation for the *ipt29* phenotype, since it can act in the root to regulate growth and development in crosstalk with CKs. IAA exhibited higher abundance compared to wild type in the short-root *ipt29* mutant (Figure 4A). This correlated with increases of IAA inactive forms such as oxIAA, oxIAA-glc, and IAA-Glu (Figure 4B). The link between auxin metabolism and the shorter root phenotype was further examined in the *ipt2* and *ipt9* single mutants. The short-root phenotype and altered IAA levels of *ipt29* were maintained only by the *ipt9* mutation and not by *ipt2* which showed wild type primary root growth and IAA levels (Figure 4 and 5). The concentration of *c*ZR and *c*Z in the roots of *ipt2* were reduced as severely as in *ipt29* compared to Col-0, while the depletion was much less pronounced in *ipt9* roots (Figure 5C). This result in combination with the phenotype of the root confirms our previous finding that the short root phenotype of *ipt9* and *ipt29* are not dependent on *c*Z concentration. In contrast, IAA content was elevated exclusively in the plants with short-root phenotypes (Figure 4) confirming the link between auxin and the phenotype in question.

We further carefully evaluated the connexion between lesions in *IPT9* and the root phenotypes. After confirming that there were no additional insertions in the original *ipt9* mutant, we isolated a new allele for this gene showing very similar phenotypic traits. Complementation analyses confirmed the connection between mutations in *IPT9* and the observed phenotypes, further supported by transgenic plants expressing a wild-type version of *IPT9* in both mutant backgrounds. Intriguingly, many of the independent overexpressor lines had roots even longer than those of Col-0. Further research is required to elucidate the potential quantitative effects of *IPT9* transcript levels on root length. Overall, our work indicates that the *IPT9* gene is essential for primary root growth. Although this gene is involved in biosynthesis of *c*Z-CKs, we hypothesize that IPT9 could be functioning through manipulation of local auxin concentration in order to control root growth.

## Acknowledgements

We thank NASC for providing the plant lines used in this work and Roger Granbom (UPSC, Umeå, Sweden) for technical assistance. The Microscopy Platform of UPSC, the Swedish Metabolomics Centre and Plant Sciences Core Facility of CEITEC Masaryk University are acknowledged for their technical support. The work was funded by the European Molecular Biology Organization (EMBO short term fellowship, project 7034) (MP). KL acknowledge Sweden’s Innovation Agency (Vinnova 2016-00504), the Knut and Alice Wallenberg Foundation (KAW 2016.0341 and KAW 2016.0352), the Swedish Research Council (VR 2018-04235) and Kempestiftelserna (JCK-2711 and JCK-1811). The Ministry of Education, Youth and Sports of the Czech Republic founded the work via the European Regional Development Fund-Project ‘Plants as a tool for sustainable global development’ (CZ.02.1.01/0.0/0.0/16_019/0000827) (ON, KD, AP, MK) and project “MSCAfellow@MUNI” [CZ.02.2.69/0.0/0.0/17_050/0008496] (MP).

## Author contributions

I.A., O.N. and M.P. conceived the project; I.A., M.A. and M.P. performed the phenotypic, treatment and confocal experiments; O.N., A.P. I.A., M.P, F.B., M.K., K.D., E. M.-B. and A.A. conducted the purification and quantification of auxins and cytokinins; E. M.-B. performed the genetic analysis, cloning and phenotyping; M.-G.V., I.A. and C.T. discussed and performed the grafting experiment. I.A., E. M.-B., M.P. and O.N. analysed and interpreted the data; I.A. and E. M.-B. made the Figures; I.A. prepared the manuscript draft; I.A., E. M.-B., and K.L. wrote the paper with input from all authors.

## SUPPORTING INFORMATION

**Supplementary Figure 1**: 24 h treatment with different CKs inhibit root growth and trigger CK signalling reporters *TCSn:GFP* and *ARR5_pro_:GUS*.

**Supplementary Figure 2**: Insertion number analysis performed in *ipt9-1* and shoot phenotypes of the *ipt9* mutants.

**Supplementary Figure 3**: Root phenotypic effects of the *35S_pro_:IPT9* transgene in mutant and wild-type backgrounds.

**Supplementary Table 1**: Primer sets used in this work

