## Supplemental Figures for "*IPT9*, a cis-zeatin cytokinin biosynthesis gene, promotes root growth"

Supporting information

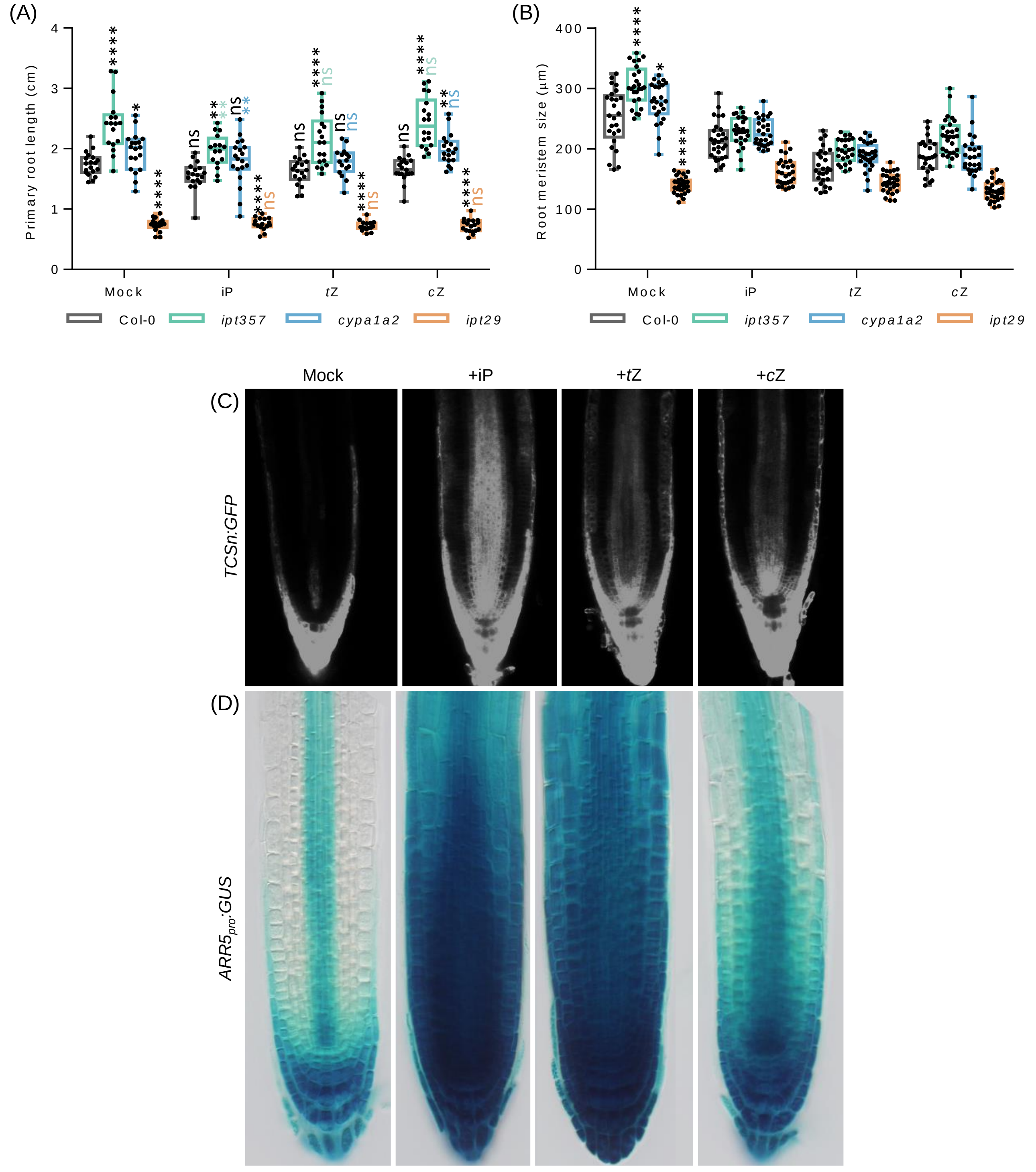


**Supplementary Figure 1**. 24 h treatment with different CKs inhibit root growth and trigger CK signalling reporters *TCSn:GFP* and *ARR5$_{pro}$:GUS*. (A, B) Primary root length (A) and root meristem (B) of the wild-type Col-0, and the *ipt357*, *cypa1a2*, and *ipt29* multiple mutants grown for 6 days in standard MS media and 24 h in MS supplemented with 100 nM of IP, tZ and cZ. Black asterisks indicate values significantly different from Col-0 mock treatment and colour asterisks indicate significantly differences from the corresponding mock genotype in a One-way ANOVA test (*p<0.05, **p<0.01, ***p<0.001, ****p<0.0001; n ≥ 16). (C, D) Cytokinin signalling reporters *TCSn:GFP* (C) and *ARR5$_{pro}$:GUS* (D) signal after 24 h of treatment with different cytokinin species.

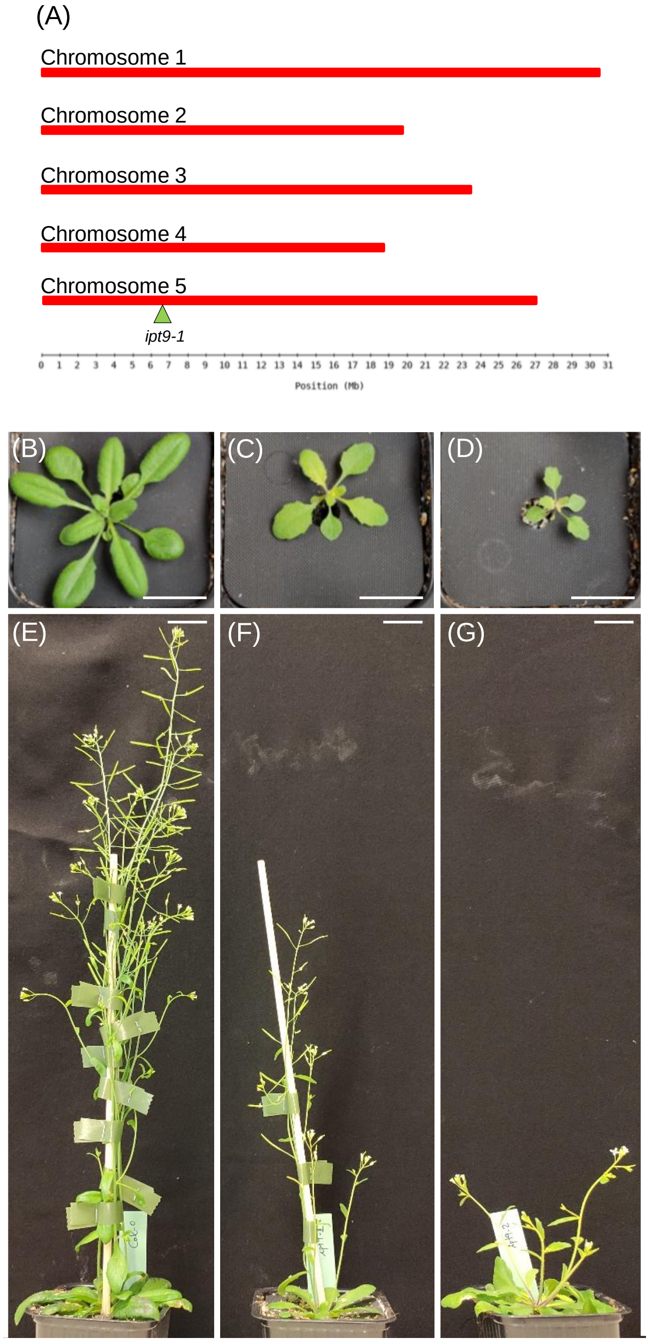


**Supplementary Figure 2**. **Insertion number analysis performed in *ipt9-1* and shoot phenotypes of the *ipt9* mutants**. (A-C) Primary root length quantification of independent transgenic families in the (A) *ipt9-1*, (B) *ipt9-2*, and (C) wild-type Col-0 background expressing *IPT9* under the constitutive 35S promoter. Different letters indicate values significantly different ($p$ < 0.05; n of each population is indicated above the genotype) in a Tukey's post-doc test. (D, E) Primary root phenotype of transgenic $35S_{pro}$*:IPT9* in the (D) *ipt9-2*, and (E) Col-0 background. (F-K) Shoot phenotype of the (F, I) wild-type Col-0 and the mutant (G, J) *ipt9-1*, and (H, K) *ipt9-2*. (L) Map of the five Arabidopsis chromosomes with indication of the position of the insertion found in *ipt9-1* (green triangle) using a tagged-sequence mapping strategy. Pictures were taken at (D, E) 7, (F-H) 23, and (I-K) 42 days. Scale bars indicate (D, E) 1 cm, and (F-K) 2 cm.

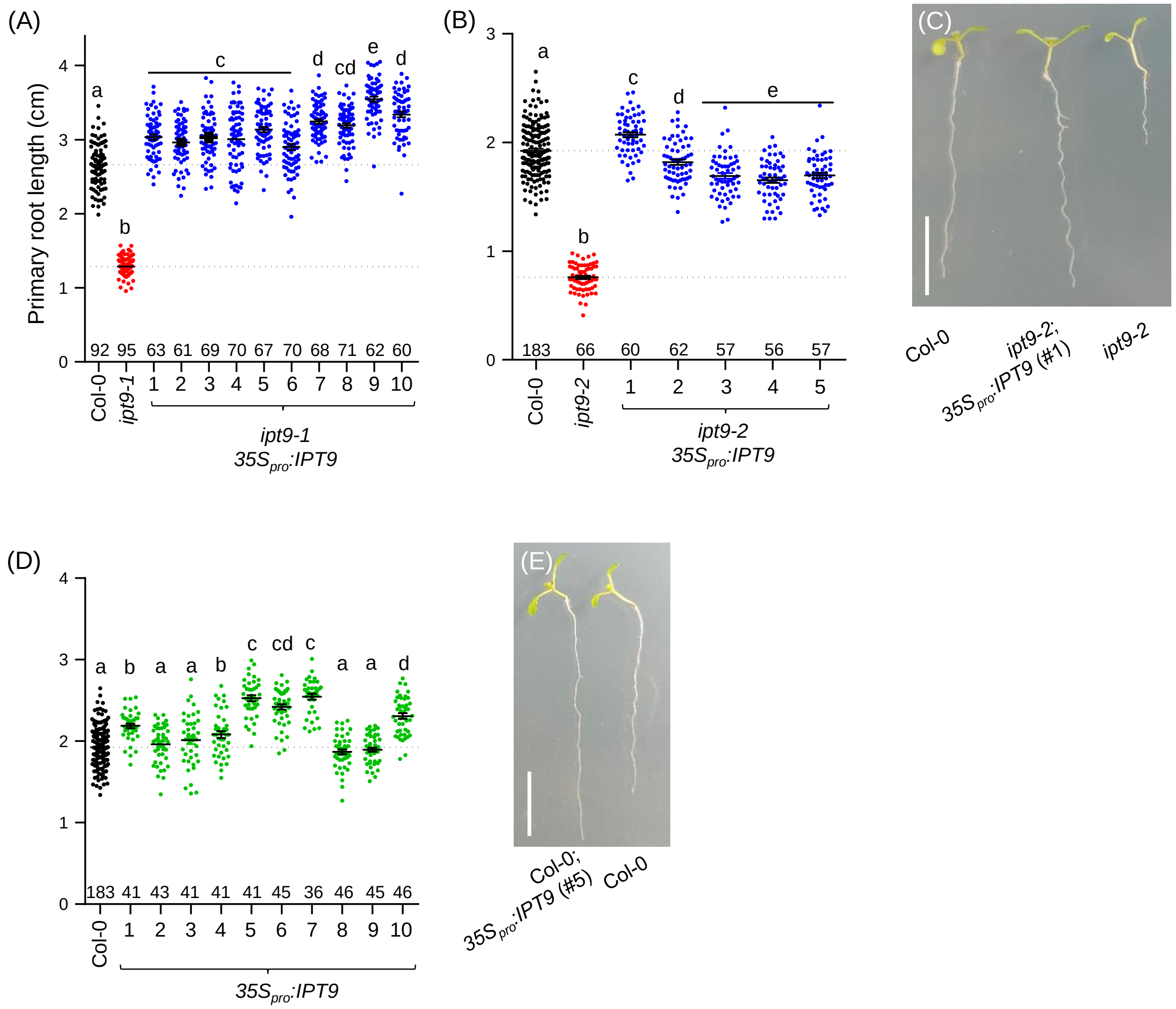


**Supplementary Figure 3. Root phenotypic effects of the 35Spro:IPT9 transgene in mutant and wild-type backgrounds**. (A, B, D) Primary root length quantification of independent transgenic families in the 7 days old (A) *ipt9-1*, (B) *ipt9-2*, and (D) wild-type Col-0 background expressing *IPT9* under the constitutive 35S promoter. Different letters indicate values significantly different ($p$ < 0.05; n of each population is indicated above the genotype) in a Tukey's post-doc test. (C, E) Primary root phenotype of transgenic *35S$_{pro}$:IPT9* in the (C) *ipt9-2*, and (E) Col-0 background. (F-K) Shoot phenotype of the (F, I) wild-type Col-0 and the mutant (G, J) *ipt9-1*, and (H, K) *ipt9-2*. (L) Map of the five Arabidopsis chromosomes with indication of the position of the insertion found in *ipt9-1* (green triangle) using a tagged-sequence mapping strategy. Scale bars indicate (C, E) 1 cm.
